# Engineering of methionine-auxotroph *Escherichia coli* via parallel evolution of two enzymes from *Corynebacterium glutamicum’s* direct-sulfurylation pathway enables its recovery in minimal medium

**DOI:** 10.1101/2024.01.08.574620

**Authors:** Matan Gabay, Inbar Stern, Nadya Gruzdev, Adi Cohen, Lucia Adriana Lifshits, Tamar Ansbacher, Itamar Yadid, Maayan Gal

## Abstract

Methionine biosynthesis relies on the sequential catalysis of multiple enzymes. *Escherichia coli*, the main bacteria used in research and industry for protein production and engineering, utilizes the three-step trans-sulfurylation pathway catalyzed by L-homoserine O-succinyl transferase, cystathionine gamma synthase and cystathionine beta lyase to convert L-homoserine to L-homocysteine. However, most bacteria employ the two-step direct-sulfurylation pathway involving L-homoserine O-acetyltransferases and O-acetyl homoserine sulfhydrylase. We previously showed that a methionine-auxotroph *E. coli* strain (MG1655) with deletion of metA, encoding for L-homoserine O-succinyl transferase, and metB, encoding for cystathionine gamma synthase, could be complemented by introducing the genes metX, encoding for L-homoserine O-acetyltransferases and metY, encoding for O-acetyl homoserine sulfhydrylase, from various sources, thus altering the *Escherichia coli* methionine biosynthesis metabolic pathway to direct-sulfurylation. However, introducing metX and metY from *Corynebacterium glutamicum* failed to complement methionine auxotrophy. Herein, we generated a randomized genetic library based on the metX and metY of *Corynebacterium glutamicum* and transformed it into a methionine-auxotrophic *E. coli* strain lacking the metA and metB genes. Through multiple enrichment cycles, we successfully isolated active clones capable of growing in M9 minimal media without external methionine supplementation. The dominant metX mutations in the evolved methionine-autotrophs *Escherichia coli* were L315P and H46R. Interestingly, we found that a metY gene encoding only the N-terminus 106 out of 438 amino acids of the wild-type MetY enzyme is functional and supports the growth of the methionine auxotroph. Recloning the new genes into the original plasmid and transforming them to methionine auxotroph *Escherichia coli* validated their functionality. These results show that directed enzyme-evolution enables the fast engineering of new active variants within the *Escherichia coli* methionine direct-sulfurylation pathway, leading to efficient complementation.

## 1 Introduction

Methionine biosynthesis is an important metabolic pathway in microorganisms and plants, encompassing a series of consecutive enzymatic steps [1–3]. As an essential amino acid for vertebrates, methionine is also highly sought for various biotechnological applications, including in the food, feed, and pharmaceutical industries [4]. Within the cellular environment, methionine is vital not only for initiating protein biosynthesis but also for protein folding and stability [5]. Moreover, it plays a significant role as the starting point for various metabolic processes, including S-adenosylmethionine (SAM), a prominent methyl group donor in cell methylation reactions [5–7].

Bacterial methionine biosynthesis relies on two distinct pathways, direct-sulfurylation and trans-sulfurylation, both convert L-homoserine to L-homocysteine, which is further converted by the enzyme methionine synthase (metE/H) to L-methionine [8–10]. However, the two pathways differ in the sulfur source used and the number of steps required to synthesize L-homocysteine [9,11,12]. In the trans-sulfurylation pathway, three steps are catalyzed by the enzymes L-homoserine O-succinyl transferase (HST; EC 2.3.1.46, MetA), cystathionine gamma synthase (CgS; EC 2.5.1.48, MetB) and cystathionine beta lyase (CbL; EC 4.4.1.13, MetC) [13]. MetA synthesizes O-succinyl L-homoserine from homoserine and succinyl CoA. MetB synthesizes cystathionine from cysteine and O-succinyl L-homoserine. MetC converts cystathionine into L-homocysteine. The direct-sulfurylation is a more efficient two-step pathway catalyzed by the enzymes L-homoserine O-acetyltransferases (HAT; EC 2.3.1.31, MetX) synthesizes O-acetyl L-homoserine from homoserine and acetyl-CoA and the O-acetyl homoserine sulfhydrylase (OAHS; EC 2.5.1.49, MetY), synthesizes L-homocysteine from O-acetyl L-homoserine and an inorganic sulfur source [8].

Most bacteria predominantly use the direct-sulfurylation pathway for methionine biosynthesis. However, *E. coli* is among the group of bacteria that employs the trans-sulfurylation pathway [14–17]. This pathway in *E. coli* is well characterized and was found to be tightly regulated due to the high energetic cost associated with methionine production [18], and with feedback inhibition from methionine and SAM [8,10,19]. Recent studies have demonstrated that introducing direct-sulfurylation genes from various organisms into *E. coli* can significantly enhance its methionine biosynthesis levels, bypassing the strict regulation of the natural trans-sulfurylation pathway [20–23]. Moreover, we recently showed that by completely removing the trans-sulfurylation enzymes encoded by metA and metB from *E. coli* and substituting them with the direct-sulfurylation genes metX and metY, *E. coli* could successfully synthesize methionine via the direct-sulfurylation pathway [24]. Interestingly, while MetX and MetY enzymes from bacteria such as *Deinococcus geothermalis* (DG) and *Cyclobacterium marinum* (CM) effectively restored methionine production in methionine-auxotrophic *E. coli*, the insertion of this pair of genes from *Corynebacterium glutamicum* (CG), a bacteria known to utilize both of the trans and direct-sulfurylation pathways, failed to do so [24,25]. To understand the molecular basis of the complementation and restore methionine biosynthesis in auxotrophic *E. coli*, we evolved the non-complementing metX and metY genes from CG to produce a functionally active enzyme pair. The concept of directed enzyme evolution involves creating a diverse genetic library and applying appropriate selection pressure, enriching and identifying variants with the desired activity profile [26– 28]. The process relies on two central steps repeated iteratively: introducing genetic diversity, either randomly or through rational design, and systematically selecting variants that display the targeted phenotypes [28,29]. In this study, we employed error-prone PCR to create a genetic library with random mutations in the MetX and MetY enzymes [30]. Following several selection rounds, we successfully identified and further characterized promising new variants of the MetX and MetY enzymes that enabled the recovery of methionine-auxotroph *E. coli* on a methionine-depleted minimal medium.

## 2 Materials and Methods

### 2.1 Generation of auxotroph *E. coli*

The generation of methionine auxotroph *E. coli* was described elsewhere [24]. Briefly, genes in *E. coli* MG1655 were deleted by the lambda red recombinase procedure [31], with the pKD4 plasmid carrying the Kn^R^ cassette as a PCR reaction template. Following each knockout cycle, the cassette was removed, and the triple knockout Δ*met*ABJ was further generated.

### 2.2 M9 minimal media

The M9 minimal media was prepared by dissolving the following components in 1L H_2_O, 6.0 g Na_2_HPO_4_, 3.0 g KH_2_PO_4_, 0.5 g NaCl, 1.5 g NH_4_Cl, 4.0 g Glucose, 0.3 g MgSO_4_, 0.02 g CaCl_2_, 250 μL trace metal mix and 250 μL Vitamin mix (Tables S2 and S3, respectively).

### 2.3 Complementation of Δ*met*ABJ with a plasmid carrying the *metX/Y* genes

Electrocompetent Δ*metABJ* mutants were transformed with pTrcHis plasmid carrying the CG *metXY* synthetic operon. Following transformation, several colonies growing on LB agar plates supplemented with 50 µg/ml ampicillin and kanamycin were tested for the presence of the correct plasmid by colony PCR (primers are listed in Table S4). Positive clones were inoculated in a 5 ml M9 and incubated overnight at 37°C (constant orbital shaking 250 rpm).

### 2.4 Growth evaluation of the bacteria with and without methionine

For the growth evaluation assays, bacteria were inoculated into a 96-well plate filled with M9 media with or without the addition of 20 μM methionine. Each well was supplemented with 100 μg/mL ampicillin and 50 μg/mL kanamycin. Growth was monitored at 37 °C with orbital shaking and absorbance at 600 nm was measured every 10 minutes in a plate reader (Synergy H1 microplate reader, BioTek).

### 2.5 Construction of a randomized genetic library of metX and metY and enrichment cycles

A genetic library was constructed based on the metX and metY genes with the GeneMorph II Random Mutagenesis Kit (Agilent, Santa Clara, CA, USA), adjusted to produce an average of 4 nonsynonymous mutations per gene. Following mutagenesis PCR, libraries were cloned back into the original vector and transformed into Δ*met*ABJ *E. coli* cells and then plated on an LB plate supplemented with 100 μg/mL amp and 50 μg/mL kana. All colonies were collected and grown for enrichment in 200 ml M9 minimal media supplemented with 100 μg/mL amp, 50 μg/mL kan, and 0.1 mM IPTG at 37 °C. For enrichment, at OD=0.8, 1 ml of the culture was rediluted in M9 and regrown. Primers used for error-prone PCR and sequencing of the construct are listed in Table S4. The full sequence of the metX/Y construct is listed in Table S5.

### 2.6 Structural modeling and visualization

Protein structures were predicted by AlphaFold2 with MMseqs2. Structural visualization was performed using UCSF ChimeraX[32].

#### 3 Results

### 3.1 *E. coli* ΔmetABJ transformed with metX/Y from CG is a methionine-auxotroph strain

To test the ability of *E. coli* to produce sufficient amounts of methionine to support growth, we used a strain auxotrophic for methionine (ΔmetABJ) expressing the CG wild-type *metX* and *metY* genes from a plasmid (ΔmetABJ-CG) [24]. We then evaluated the ability of this strain to grow on M9 minimal media, which contained glucose, ammonium chloride, and MgSO_4_ as the carbon, nitrogen, and sole sulfate sources, respectively. As depicted in Figure 1, the ΔmetABJ-CG strain showed growth in the presence of methionine (red curve) but failed to grow without it (blue curve). This indicates that methionine limits bacterial growth and that the metX and metY genes from CG are insufficient for autonomous methionine production in *E. coli*, thus necessitating external supplementation of methionine.

**Figure 1.**
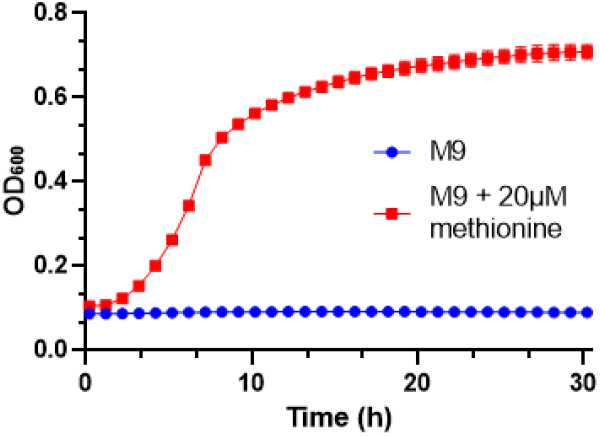
Methionine Dependency of the ΔmetABJ-CG Strain. The graph shows the growth dependency of the ΔmetABJ-CG strain on methionine in M9 minimal media, comparing the growth curve of the strain supplemented with 20 µM methionine (red curve) against its growth without methionine supplementation (blue curve). Data points represent the mean ± standard deviation (SD) from three independent replicates.

### 3.2 Identifying active MetX/Y variants in a methionine auxotrophic *E. coli*

To identify metX and metY variants from CG capable of supporting methionine synthesis in the *E. coli* ΔmetABJ, we generated a randomized genetic library from these genes. We calibrated the error rate of the PCR such that approximately four non-synonymous mutations will be inserted per 1000 base pairs. The actual library size, resulting from the transformation efficiency, was ∼1.5 × 10^5^. This library was then introduced into ΔmetABJ *E. coli*, initiating the selection process to identify active variants of the MetX/Y enzymes. We initiated this process by transforming these cells with our randomized library and cultivating them in the M9 minimal medium. Figure 2A illustrates the enrichment process used to select variants that effectively complement ΔmetABJ *E. coli* cells, which was performed by diluting the primary culture by a factor of 1000 upon reaching an OD_600_=0.8, allowing for regrowth in the M9 medium. With each cycle corresponding to about 10 generations (0.008 X 2^10^), this dilution and regrowth step was repeated for five cycles. After each cycle, we plated aliquots from the culture on M9 agar plates containing ampicillin and kanamycin for selection and confirmed their ability to grow in the absence of methionine. Moreover, the presence of metY and metX genes was verified using colony PCR. Figure 2B illustrates the growth curves of these enrichment cycles. Notably, advanced cycles exhibited an accelerated growth rate, indicating the successful enrichment of methionine-producing bacterial variants.

**Figure 2.**
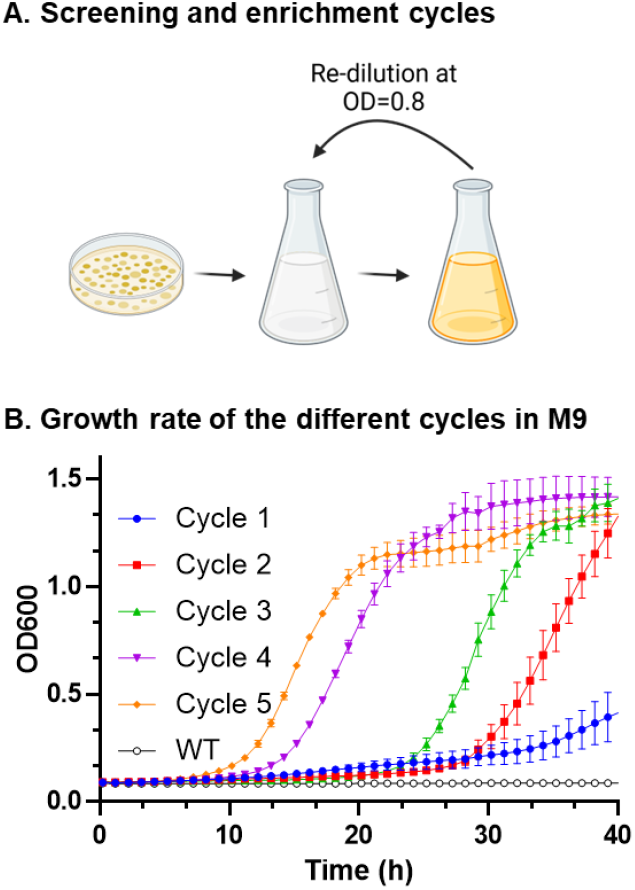
Enrichment of Active MetX/Y Variants. **(A)** Schematic of the enrichment process: The panel illustrates the sequential steps to select active genetic variants from the library. **(B)** The growth patterns of the wild-type strain and enrichment cycles following the generation of the genetic library demonstrate the enhancement in growth rates with each cycle.

### 3.3 Sequence and structural analysis of evolved MetX and MetY variants

Following multiple rounds of enrichment, we sequenced multiple colonies from the fifth cycle (C5) and characterized the mutations. Sequencing results showed two dominant mutations in metX (MetX-C5) identified in all tested colonies and a significant deletion in metY in part of the colonies. The enzyme amino acid sequences of both the wild-type and variants are presented in Table S1. Figure 3A highlights the mutations in MetX-C5, specifically an arginine to histidine substitution at position 46 (R46H) and a leucine to proline substitution at position 315 (L315P). Figure 3B shows a sequence alignment, comparing the wild-type MetY gene with its truncated variant (MetY-del). This shortened variant comprises only the first 106 amino acids, a substantial reduction from the 438 amino acids of the full-length wild-type enzyme.

**Figure 3.**
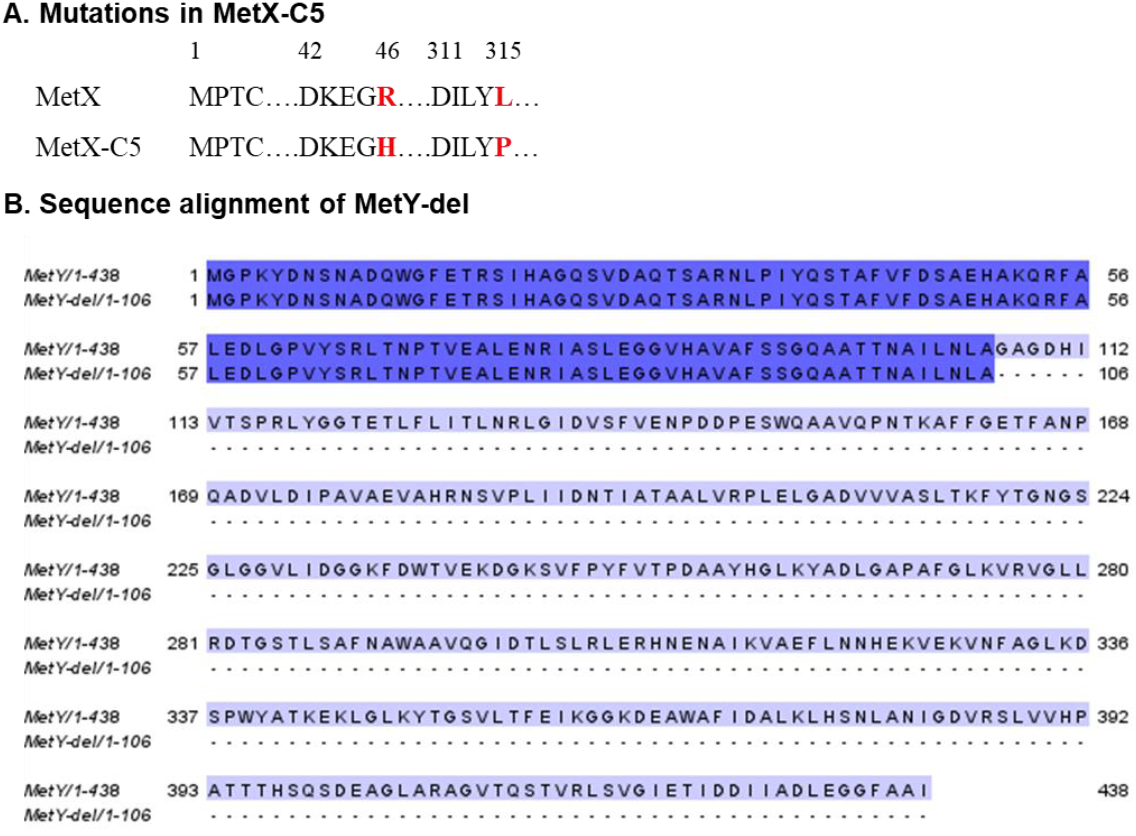
Sequence and mutations in the evolved MetX-C5 and MetY-del Variants. **(A)** Specific single-point mutations of R46H and L315P were discovered in the MetX enzyme and are highlighted in red. **(B)** Alignment of the sequence of MetY-del with the wild-type enzyme. The 106 amino acids retained in the MetY-del variant are highlighted in dark purple.

### 3.4 Analysis of MetY-del variant

To decipher the functionality of the truncated MetY-del, we first employed BLAST to find similar protein sequences. This revealed a protein from *Corynebacterium diphtheriae*, annotated as O-acetylhomoserine amino carboxypropyl transferase (E.C. 2.5.1.49, NCBI reference sequence CAB0498630.1), which shares considerable similarity with MetY-del in both size and sequence (Fig. 4A). Complementing the sequence analysis, we utilized AlphaFold to model the structures of both proteins. Figure 4B displays the structural alignment of MetY-del (in light blue) and the O-acetylhomoserine amino carboxypropyl transferase (in green). In agreement with the sequence similarity, the structural alignment revealed considerable structural similarity, except for differences in the α-helix regions. This supports the notion that deleting the large amino acids segment resulting in MetY-del can indeed generate a functional enzyme.

**Figure 4.**
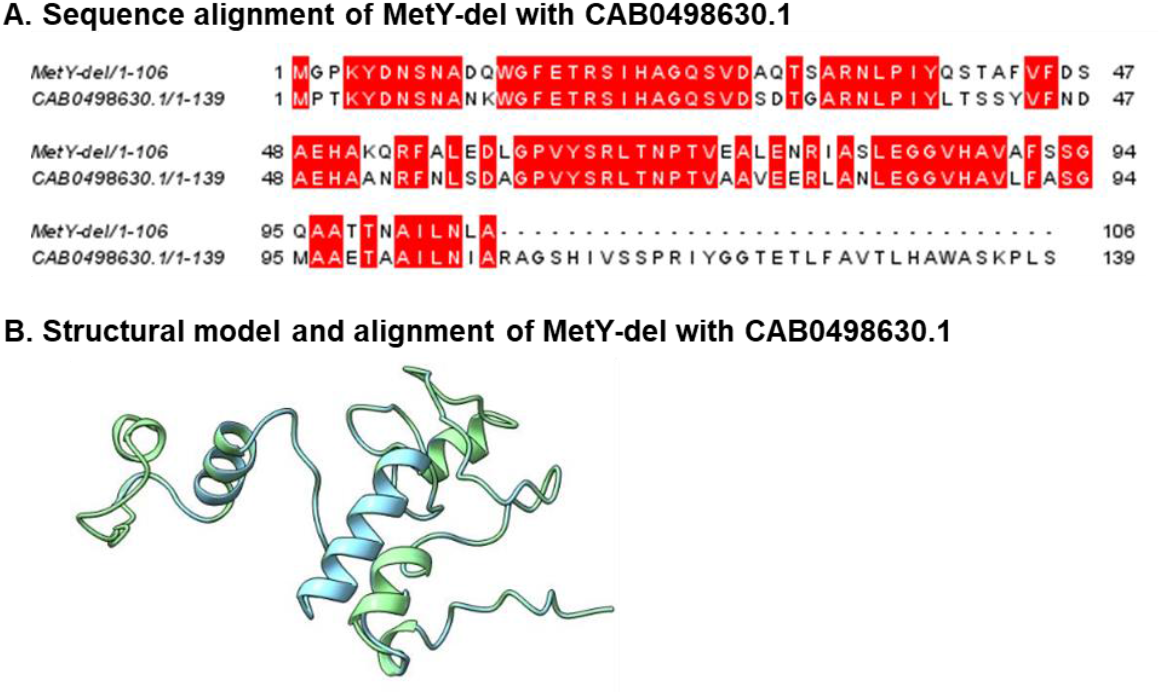
Analysis of MetY-del and O-Acetylhomoserine amino carboxypropyl transferase from *C. diphtheriae*. **(A)** Sequence alignment of the two enzymes (MetY-del in the top row). Sequence alignment was done with JalView version 2.11.3.2. **(B)** Structure alignment of the two enzymes (MetY-del in light blue).

### 3.5 Sequence alignment and structural analysis of MetX

**Figure 5A** shows the alignment of MetX of CG with additional MetX enzymes. The blue arrow points to the position in which the two mutations occurred. Interestingly, the proline residue acquired by the CG enzyme following the L315P mutation is highly conserved in other MetX enzymes. This segment is central to the catalytic site. Figure 5B shows the 3D model of MetX CG enzyme, with the mutated residues His and Pro highlighted in red on top of the wild-type enzyme (gray). As shown in other MetX enzyme structures, the acquisition of the conserved proline residue supports the correct structural formation of the catalytic site tunnel and architecture of the conserved catalytic triad [33]. Indeed, as in other bacterial species, a Pro in position adjacent to the catalytic site is indeed conserved. In MetX-C5, the Pro residue is positioned at the N-terminus of an alpha helix, composed of residues Y316 to N325 following L313 and Y314, both are part of the acetyl-moiety binding site[34]. The Pro single rotatable bond and its ability to break the helix continuation makes the L315P at the beginning of the Y316 - N325 α helix contribute to the overall stabilization of the protein. An additional contribution could arise from specific interactions involving the pyrrolidine ring atoms [35]. Here, the P315 ring could bind with the adjacent backbone oxygen of Asp311, one of the three amino acids of the catalytic triad. Such stabilization could contribute to the enhanced catalytic activity of the enzyme in the cell. The R46H mutation most likely did not change the enzyme activity. Considering that the Arg is buried within a positive pocket formed by Glu44 and Asp42, a mutation to another positive, yet less flexible residue as His is not expected to induce a crucial stabilizing effect. However, the overall contribution of both mutations is positive.

**Figure 5.**
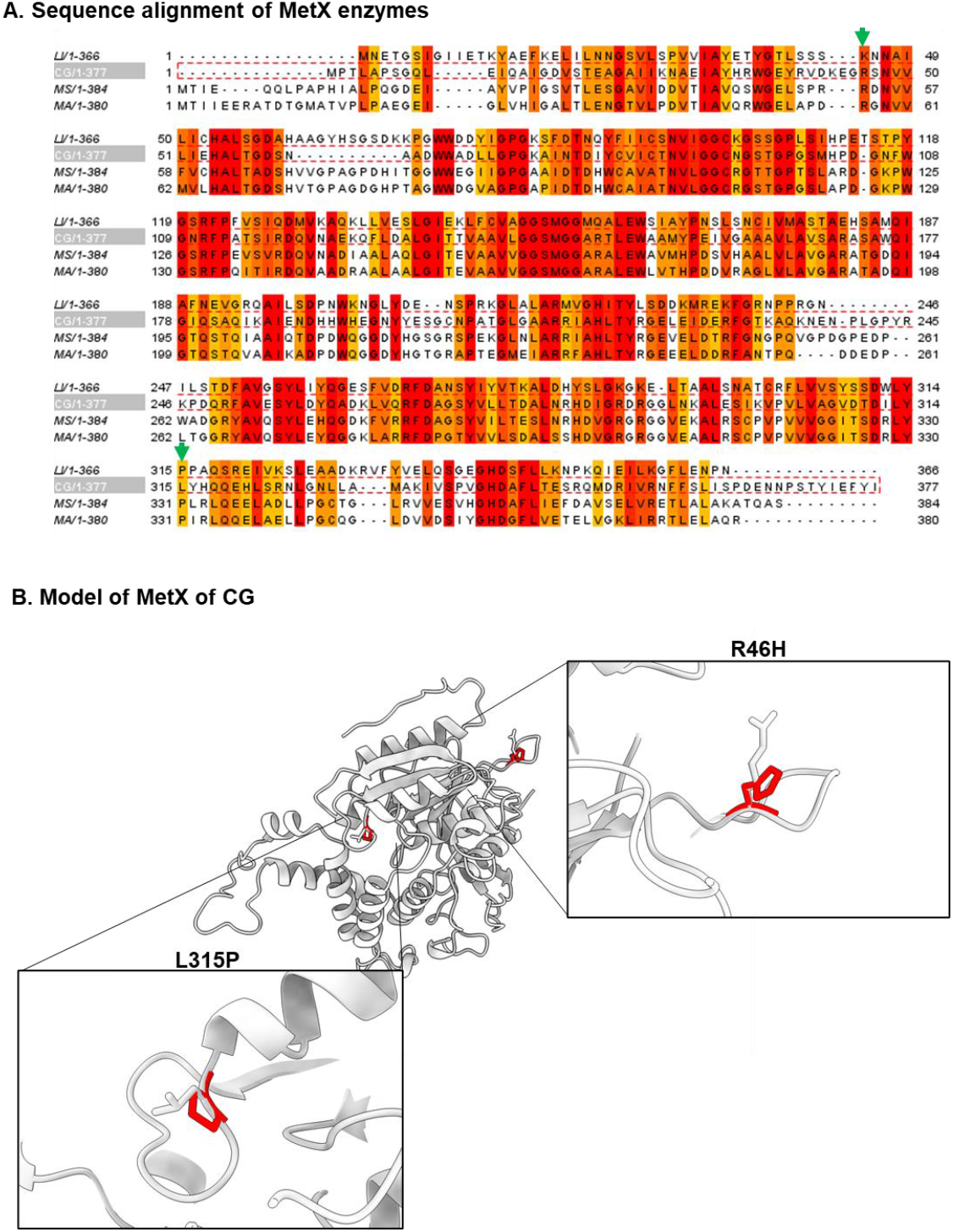
Analysis of MetX and MetX-C5. **(A)** Sequence alignment of MetX CG with MetX of *Leptospira interrogans* (LI), *Mycolicibacterium smegmatis* (MS), and *Mycobacteroides abscessus* (MA). Green arrows show the position of the R46H and L315P mutations. Sequence alignment was done with JalView version 2.11.3.2. **(B)** The structure of wild-type MetX CG and the MetX-C5 with the two mutations are shown in red color.

### 3.6 Validating the activity of the new MetX and MetY Variants

To confirm the functionality of the new MetX and MetY variants, each variant gene was re-cloned into the original empty plasmid and then used to transform the original *E. coli* ΔmetABJ strain. This approach ensures that unintended mutations elsewhere in the plasmid or in the bacterial genome do not influence any observed activity. It also imparts uniform regulatory conditions for all genetic configurations. **Figure 6** displays the growth curves in M9 media of *E. coli* ΔmetABJ harboring each modified plasmid. Notably, the plasmids ISM5 and ISM7, carrying wild-type MetY and MetY-del, respectively, along with the MetX-C5 variant, showed successful bacterial proliferation. The strain with wild-type MetY exhibited a higher growth rate, suggesting that the activity of the full-length protein is better than that of the shorter variant MetY-del. In contrast, strains with other genetic configurations failed to grow, thereby validating the functionality of these specific enzyme variants and underscoring the necessity of both enzymes for bacterial growth.

**Figure 6.**
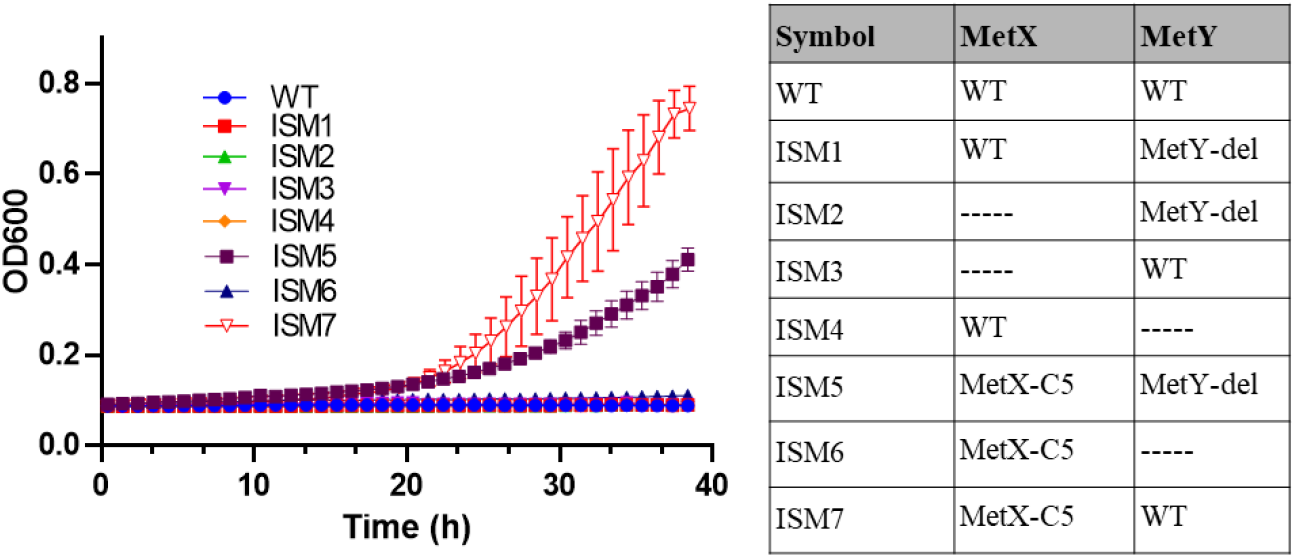
Comparative growth analysis of *E. coli* ΔmetABJ with various MetX/Y variants. Each growth curve of *E. coli* ΔmetABJ represents a different combination of MetX and MetY variants within the ISM plasmids.

## 4 Discussion

Methionine is a critical amino acid in food, feed, and pharmaceutical industries, and its efficient biosynthesis is of significant public and commercial interest. The potential of engineering *E. coli*, a workhorse in industrial biotechnology, to enhance methionine production is particularly promising. By altering its natural methionine biosynthesis pathway through the incorporation of evolved MetX and MetY enzymes, one can potentially boost the production of this essential amino acid. This will not only address the need for methionine but will also open avenues for more sustainable and cost-effective production methods. The ability to engineer *E. coli* for enhanced methionine biosynthesis is thus a critical step forward for such biotechnological innovations. This study focused on the evolution of the direct-sulfurylation enzymes MetX and MetY from *C. glutamicum* in a methionine-auxotroph *E. coli*. Although *E. coli* is a common bacteria for the production of amino acids, its methionine biosynthesis pathway relies on the less abundant and heavily regulated trans-sulfurylation pathway, a challenge for efficient methionine production[36,37]. Previous studies have shown that clearance of negative regulation of the methionine biosynthesis routes has enhanced the ability to produce methionine [37–40]. Herein, we explored the ability to harness laboratory evolution of MetX and MetY, to transform an auxotroph-methionine *E. coli* into a prototroph. The evolution of the CG enzymes MetX and MetY led to the identification of specific mutations that conferred functionality in the introduced direct-sulfurylation enzymes in *E. coli*. The mutations identified in MetX and MetY, particularly those in the active site of MetX, likely improve the enzymes’ catalytic efficiency or substrate specificity to fit to the *E. coli* cellular environment or evade the cellular regulation mechanism within the cell. Indeed, the deletion found in MetY points towards a potential structural simplification that may be more compatible with *E. coli*’s cellular machinery. Despite the deletion of a large segment, MetY-del retained the ability to support the complementation of the methionine auxotroph. This result is congruent with the evolutionary notion that proteins can undergo truncation while maintaining essential functional characteristics [41,42], often leading to increased efficiency or altered regulatory control. An additional point within the context of the functional validation of the evolved enzymes in the ΔmetABJ *E. coli* is the adaptability of metabolic pathways. Herein, we simultaneously evolved two enzymes within the same pathway on the same plasmid. An interesting point to consider is whether such parallel evolution is important. On the one hand, the mutational space is dramatically increased due to the larger operon. However, if cooperative activity of the two enzymes is beneficial, this enables the screening for sequence combinations that otherwise could be neglected.

A question arises regarding the most efficient evolutionary path to generate functional enzymes. We relied on introducing random mutations throughout the gene, mimicking the stochastic nature of evolutionary changes. This method is useful when multiple enzymes catalyze a series of chemical reactions, as it allows for exploring a vast and often unexplored sequence space. Rational design, on the other hand, involves introducing specific mutations based on known structures, mechanisms, interactions, and known metabolic flux. While this method can be more targeted, it is limited by the number of calculated mutations and potential combinations and may miss beneficial mutations that are not intuitively obvious. An additional important factor is the evolutionary starting point. Starting with a non-active enzyme, as in the case of the CG-derived MetX and MetY in *E. coli*, can be challenging due to the initial lack of function. However, it offers the opportunity to discover novel adaptations and functionalities that may not be present in the native enzyme. This approach can yield enzymes better suited to new environments or substrates. On the other hand, beginning with an active enzyme allows for a more incremental improvement approach, focusing on enhancing or altering specific properties such as substrate specificity, stability, or catalytic efficiency. While this approach may offer a quicker path to improved function, it may not cover the large space of functional diversity achievable when starting from a non-active baseline. Of note, we sequenced representative colonies of the final enrichment cycles and found a dominant mutation in Metx-C5 and the single variant MetY-del. However, other variants in earlier enrichment cycles could exhibit additional variants capable of producing methionine in the auxotroph bacteria. Although less efficient than MetX-C5, such variants could set a starting point for additional genetic libraries, leading to new and unexplored sequences. The search for alternative sequences, together with structure-function studies of current variants, is the focus of our following studies.

## 5 Conclusions

Our study shows that through the substitution of the native trans-sulfurylation *E. coli* enzymes with direct-sulfurylation and by the application of directed enzyme evolution, we engineered *E. coli* capable of producing a sufficient amount of methionine to support growth. Together with improving regulation and metabolic flux, enzyme optimization could be a promising approach to support methionine biosynthesis and thus pose an important avenue for a broad range of biotechnological applications.

## Acknowledgments

This research was supported by the Israeli Ministry of Agriculture (grant# 21-36-0003), the Gertner Institute for Medical Nano System at Tel Aviv University and the Israeli Ministry of Science and Technology (grant# 0005749). L.A.L is grateful for Ph.D. scholarship financial support from the ADAMA Center for Novel Delivery Systems and the Gertner Institute for the Medical Nano System in TAU.

## Author contributions

M.G, I.S and N.G performed the experimental research and analysis. A.C and L.A.L assisted with experiments and data analysis. T.A. analyzed the data and wrote the manuscript. I.Y, and M.G. analyzed the data, conceived and supervised the research and wrote the manuscript. All authors reviewed and approved the manuscript.

