## Supplementary material for "Engineering of methionine-auxotroph *Escherichia coli* via parallel evolution of two enzymes from *Corynebacterium glutamicum’s* direct-sulfurylation pathway enables its recovery in minimal medium": Supp

**Table S1. Protein sequences.**

| Protein name | Sequence |
|---|---|
| MetX CG | MPTLAPSGQLEIQAIGDVSTEAGAIIKNAEIAYHRWGEYRVDKEGRSNVVLIEHALTGDSN<br>AADWWADLLGPGKAINTDIYCVICTNVIGGCNGSTGPGSMHPDGNFWGNRFPATSIRD<br>QVNAEKQFLDALGITTVAAVLGGSMGGARTLEWAAMYPEIVGAAAVLAVSARASAWQIG<br>IQSAQIKAIENDHHWHEGNYYESGCNPATGLGAARRIAHLTYRGELEIDERFGTKAQKNE<br>NPLGPYRKPDQRFAVESYLDYQADKLVQRFDAGSYVLLTDALNRHDIGRDRGGLNKAL<br>ESIKVPVLVAGVDTDILYLYHQQEHLSRNLGNLLAMAKIVSPVGHDAFLTESRQMDRIVR<br>NFFSLISPDENNPSTYIEFYI |
| MetX CG R46H, L315P | MPTLAPSGQLEIQAIGDVSTEAGAIIKNAEIAYHRWGEYRVDKEGHSNVVLIEHALTGDSN<br>AADWWADLLGPGKAINTDIYCVICTNVIGGCNGSTGPGSMHPDGNFWGNRFPATSIRD<br>QVNAEKQFLDALGITTVAAVLGGSMGGARTLEWAAMYPEIVGAAAVLAVSARASAWQIG<br>IQSAQIKAIENDHHWHEGNYYESGCNPATGLGAARRIAHLTYRGELEIDERFGTKAQKNE<br>NPLGPYRKPDQRFAVESYLDYQADKLVQRFDAGSYVLLTDALNRHDIGRDRGGLNKAL<br>ESIKVPVLVAGVDTDILYPYHQQEHLSRNLGNLLAMAKIVSPVGHDAFLTESRQMDRIVR<br>NFFSLISPDENNPSTYIEFYI |
| MetY CG | MGPKYDNSNADQWGFETRSIHAGQSVDAQTSARNLPIYQSTAFVFDSAEHAKQRFALE<br>DLGPVYSRLTNPTVEALENRIASLEGGVHAVAFSSGQAATTNAILNLAGAGDHIVTSPRL<br>YGGTETLFLITLNRLGIDVSFVENPDDPESWQAAVQPNTKAFFGETFANPQADVLDIPAV<br>AEVAHRNSVPLIIDNTIATAALVRPLELGADVVVASLTKFYTGNGSGLGGVLIDGGKFDWT<br>VEKDGKSVFPYFVTPDAAYHGLKYADLGAPAFGLKVRVGLLRDTGSTLSAFNAWAAVQ<br>GIDTLSLRLERHNENAIKVAEFLNNHEKVEKVNFAGLKDSPWYATKEKLGLKYTGSVLTF<br>EIKGGKDEAWAFIDALKLHSNLANIGDVRSLVVHPATTTHSQSDEAGLARAGVTQSTVRL<br>SVGIETIDDIIADLEGGFAAI |
| MetY-del | MGPKYDNSNADQWGFETRSIHAGQSVDAQTSARNLPIYQSTAFVFDSAEHAKQRFALE<br>DLGPVYSRLTNPTVEALENRIASLEGGVHAVAFSSGQAATTNAILNLA |
| CAB0498630 | MPTKYDNSNANKWGFETRSIHAGQSVDSDTGARNLPIYLTSSYVFNDAEHAANRFNLSD<br>AGPVYSRLTNPTVAAVEERLANLEGGVHAVLFASGMAAETAAILNIARAGSHIVSSPRIYG<br>GTETLFAVTLHAWASKPLS |

**Table S2. Vitamin mix**

| Vitamin mix | 500 mL |
|---|---|
| p-aminobenzoic acid | 0.5 gr |
| Niacin | 0.5 gr |
| Pyridoxine | 0.5 gr |
| Riboflavin | 0.5 gr |
| Thiamin HCl | 0.5 gr |
| Choline HCl | 0.5 gr |
| d-Biotin | 1 mg |
| Store at 4 °C in the dark after autoclaving | |

**Table S3. Trace element solution**

| **Trace elements solution** | **250 mL (H2O** |
|---|---|
| $H_3BO_3$ | 100 mg |
| $CuSO_4$ $5H_2O$ | 100 mg |
| Iron chloride $4H_2O$ | 200 mg |
| $MnCl_2$ | 200 mg |
| $NaMoO_4$ $2H_2O$ | 200 mg |
| $ZnSO_4$ $7H_2O$ | 2 gr |
| Add HCl until the brown murky solution turns clear and yellow, filter sterile through 0.2 µM, keep in the dark. | |

**Table S4. Primers used in this study**

| Primer name | Sequence | Use |
|---|---|---|
| cTrc_Ins_Fw | ATAAGGAGGAATAAACCATGGGTCCG | Error-prone PCR and sequencing |
| cTrc_Ins_Rev | GGTACCAGCTGCAGATCTCGAGTTA | Error-prone PCR and sequencing |
| metY_F_SEQ | AAAGTTCGTGTGGGCCTGCT | Sequencing |
| metY_R_SEQ | AGCAGGCCCACACGAACTTT | Sequencing |
| metX_F_SEQ | TGGTTCTGATCGAACACGCGC | Sequencing |

**Table S5. Sequence of MetYX CG construct.**

| Gene | Sequence |
|---|---|
| Wld-type MetXY gene of CG that was cloned and transformed to the E. coli | > metYX_CG_ OQ291222<br>gaattcTTTATTCTTGACACTAGTCGGCCAAAATGATATAATACCTGAGTT<br>TAACTTTAAGAGAGGTATATATTACCATGGGTCCGAAGTACGACAACA<br>GCAACGCGGATCAGTGGGGCTTCGAGACCCGTAGCATCCACGCGGG<br>TCAGAGCGTTGACGCGCAAACCAGCGCGCGTAACCTGCCGATTTACC<br>AGAGCACCGCGTTCGTGTTTGACAGCGCGGAGCACGCGAAACAACGT<br>TTCGCGCTGGAAGATCTGGGCCCGGTTTATAGCCGTCTGACCAACCC<br>GACCGTGGAGGCGCTGGAAAACCGTATTGCGAGCCTGGAGGGTGGC<br>GTTCATGCGGTGGCGTTTAGCAGCGGTCAGGCGGCGACCACCAACG<br>CGATCCTGAACCTGGCGGGTGCGGGTGACCACATTGTTACCAGCCCG<br>CGTCTGTATGGTGGCACCGAAACCCTGTTCCTGATCACCCTGAACCGT<br>CTGGGCATTGATGTTAGCTTTGTGGAGAACCCGGATGATCCGGAAAG<br>CTGGCAGGCGGCGGTTCAACCGAACACCAAGGCGTTCTTTGGCGAGA<br>CCTTTGCGAACCCGCAAGCGGACGTGCTGGATATCCCGGCGGTTGCG<br>GAAGTGGCGCACCGTAACAGCGTTCCGCTGATCATTGACAACACCATT<br>GCGACCGCGGCGCTGGTGCGTCCGCTGGAGCTGGGTGCGGATGTGG<br>TTGTGGCGAGCCTGACCAAGTTCTACACCGGTAACGGCAGCGGTCTG<br>GGTGGCGTTCTGATCGACGGTGGCAAATTTGATTGGACCGTGGAAAA<br>GGACGGCAAAAGCGTTTTCCCGTATTTTGTTACCCCGGATGCGGCGT<br>ACCACGGTCTGAAGTATGCGGATCTGGGTGCGCCGGCGTTTGGTCTG<br>AAAGTTCGTGTGGGCCTGCTGCGTGACACCGGTAGCACCCTGAGCGC<br>GTTTAACGCGTGGGCGGCGGTTCAAGGCATCGATACCCTGAGCCTGC<br>GTCTGGAGCGTCACAACGAAAACGCGATTAAGGTGGCGGAGTTCCTG<br>AACAACCACGAGAAGGTTGAAAAAGTGAACTTTGCGGGTCTGAAGGAT<br>AGCCCGTGGTACGCGACCAAGGAAAAACTGGGCCTGAAATATACCGG<br>TAGCGTGCTGACCTTCGAGATCAAGGGTGGCAAAGACGAAGCGTGGG<br>CGTTTATTGATGCGCTGAAACTGCACAGCAACCTGGCGAACATCGGC<br>GACGTTCGTAGCCTGGTTGTGCATCCGGCGACCACCACCCATAGCCA<br>AAGCGATGAGGCGGGCCTGGCGCGTGCGGGTGTGACCCAAAGCACC<br>GTTCGTCTGAGCGTGGGTATCGAGACCATTGACGATATCATTGCGGA<br>CCTGGAAGGTGGCTTCGCGGCGATTTAAGGATCCAGAGGTATATATTA<br>atgCCGACCCTGGCGCCGAGCGGTCAGCTGGAGATCCAAGCGATTGG<br>TGACGTTAGCACCGAGGCGGGCGCGATCATTACCAACGCGGAAATTG<br>CGTACCACCGTTGGGGTGAGTATCGTGTGGACAAAGAAGGCCGTAGC<br>AACGTGGTTCTGATCGAACACGCGCTGACCGGTGATAGCAACGCGGC<br>GGACTGGTGGGCGGATCTGCTGGGTCCGGGCAAGGCGATCAACACC<br>GACATTTACTGCGTTATCTGCACCAACGTGATCGGTGGCTGCAACGG<br>CAGCACCGGTCCGGGCAGCATGCACCCGGATGGTAACTTCTGGGGC<br>AACCGTTTTCCGGCGACCAGCATTCGTGACCAGGTTAACGCGGAGAA<br>ACAATTCCTGGATGCGCTGGGTATTACCACCGTTGCGGCGGTGCTGG<br>GTGGCAGCATGGGTGGCGCGCGTACCCTGGAGTGGGCGGCGATGTA<br>TCCGGAAACCGTTGGTGCGGCGGCGGTGCTGGCGGTTAGCGCGCGT<br>GCGAGCGCGTGGCAGATCGGCATTCAGAGCGCGCAAATCAAGGCGA<br>TTGAAAACGATCACCACTGGCACGAGGGTAACTACTATGAAAGCGGCT<br>GCAACCCGGCGACCGGTCTGGGTGCGGCGCGTCGTATTGCGCACCT<br>GACCTACCGTGGTGAGCTGGAAATCGACGAGCGTTTTGGCACCAAGG<br>CGCAGAAAAACGAAAACCCGCTGGGTCCGTATCGTAAGCCGGATCAA<br>CGTTTCGCGGTTGAGAGCTACCTGGACTATCAGGCGGATAAACTGGT<br>TCAACGTTTTGACGCGGGTAGCTACGTGCTGCTGACCGATGCGCTGA |

| | |
|---|---|
| | ACCGTCACGACATTGGCCGTGATCGTGGTGGCCTGAACAAGGCGCTG<br>GAGAGCATTAAAGTGCCGGTTCTGGTGGCGGGCGTTGACACCGATAT<br>CCTGTACCCGTATCACCAGCAAGAACACCTGAGCCGTAACCTGGGTA<br>ACCTGCTGGCGATGGCGAAAATCGTTAGCCCGGTGGGTCATGATGCG<br>TTCCTGACCGAAAGCCGTCAAATGGATCGTATTGTGCGTAACTTCTTT<br>AGCCTGATCAGCCCGGACGAGGATAACCCGAGCACCTACATTGAATT<br>TTATATCTAACTCGAGCAACCTGGAGGCGGGCGCAGGCCCGCCTTTT<br>aagctt |
